# Insulin receptor signaling in gustatory cells suppresses sugar taste sensitivity and feeding in *Drosophila*

**DOI:** 10.64898/2026.01.29.702574

**Authors:** Christian Arntsen, Lindsey Earle, Macy Ingersoll, Molly Stanley

## Abstract

Dynamic sensory processing helps animals adjust behaviors in response to changing environments and internal states. Hunger is a salient internal cue that can directly modify chemosensory signaling to promote homeostatic behavior. In *Drosophila melanogaster*, hunger enhances the sensitivity of sweet gustatory receptor neurons (GRNs) to encourage feeding, and here we investigated candidate hunger/satiety hormone receptors to identify mechanisms of metabolic regulation in primary taste cells. The conserved insulin receptor (InR) is expressed in sweet GRNs, and targeted *InR* knockdown and inactivation within these cells elevated sucrose sensitivity, increased sucrose consumption, and altered feeding behavior under sated conditions. Additionally, sweet cellular responses to sucrose were reciprocally affected by inactive and overactive InR signaling. These findings reveal that InR signaling can modulate sweet sensitivity at the level of primary taste cells, suppressing feeding by reducing taste responsiveness under specific metabolic conditions.

## INTRODUCTION

Behavioral plasticity allows organisms to appropriately respond to shifting external and internal stimuli^1–5^. Hunger is a conserved internal cue that regulates widespread behaviors, including locomotion^6–8^, reproduction^9–12^ and decision making^13,14^ to prioritize feeding. A key mechanism of this hunger modulation involves direct modification of chemosensory processing within the nervous system. Food deprivation heightens both olfactory and gustatory sensitivity to certain cues in humans and other animals^15–19^. In particular, sweet taste elicits strong neural and behavioral responses across species^20–24^, making the control of sweet sensation and perception especially important for flexible feeding^25^. For example, gustatory alliesthesia^26^ describes how hunger enhances the hedonic expression of sweet stimuli to encourage feeding and maintain energy homeostasis^25^. Despite neuroimaging evidence for this phenomenon^27–29^, the precise mechanisms underlying the integration of metabolic information within gustatory circuits remain unclear.

Previous research on the interactions between taste and hunger has primarily focused on central processing^27–29^ with less known about how metabolic signals may influence peripheral taste responses. The fruit fly, *Drosophila melanogaster*, is an advantageous model organism for studying gustatory neurobiology^30^ because primary taste responses can be recorded in awake animals with intact internal states^31–38^. The robust genetic tools^39^ combined with this powerful *in vivo* imaging approach^38,40,41^ in *Drosophila* have greatly improved our understanding of taste encoding and how gustatory circuits coordinate feeding behavior^31–37,41,42^. Flies detect diverse chemical tastants using gustatory receptor neurons (GRNs) located in various structures, most prominently within taste sensilla populating the surface of the labellum - the fly’s primary peripheral taste organ^43–45^. Of the five recognized labellar GRN classes^31^, sugar-sensing “sweet” GRNs are the most abundant and are characterized by expression of nine sugar-specialized gustatory receptors (GRs)^46,47^ that function as tetrameric ligand-gated cation channels^48^. Besides sugar, these taste cells respond to many attractive chemical compounds to ultimately drive appetitive feeding responses^31,32,49–53^.

Importantly, recording from mammalian taste cells requires anesthesia, a treatment that significantly alters internal status and endocrine signaling^54,55^. In flies, sweet GRNs are amenable to *in vivo* calcium imaging in awake animals, which has revealed that starvation enhances taste sensitivity at the level of primary sweet cells^56–58^. Dopamine receptor signaling is one mechanism of this starvation-induced increase in sweet sensitivity^56,57^, and conversely, factors that suppress sweet GRN responses also alter taste sensitivity and impact feeding^34,35,59^. Efforts to find other neuromodulators in sweet GRNs have identified *CCKLR-17D3*, a receptor for the *Drosophila* cholecystokinin analogue: drosulfakinin (DSK)^60,61^, but the role of canonical hunger/satiety hormones in maintaining dynamic gustatory processing has not been fully explored.

Metabolic hormones maintain energy homeostasis through various mechanisms, including hunger/satiety balance in the brain^62–64^. In mice, glucagon signaling affects sweet taste responsiveness^65^ and mammalian type II taste receptor cells (TRCs) express insulin receptors^66^, indicating the possibility of endocrine signaling in primary gustatory cells. In flies, *Drosophila*-insulin-like-peptides (Dilps) are a group of hormones with diverse roles in metabolism and reproduction that act through a conserved insulin receptor (InR)^67^, and AKHR is a receptor for the *Drosophila* glucagon analog, adipokinetic hormone (AKH). Akin to mammals, Dilp and AKH signaling changes with metabolic status to help regulate feeding^68–75^. Furthermore, InR signaling has been found to impact olfactory sensitivity during food-deprivation^76^, but its potential function in primary gustatory cells remains unclear. Our work aims to characterize the role of hunger/satiety hormone receptors in sweet GRN modulation, sweet taste sensitivity, and overall feeding behavior under different metabolic conditions. Using single-cell transcriptomic evidence, we found that *InR* and *AKHR* were expressed in many sweet GRNs and find that InR contributes to satiety-state feeding regulation at the level of primary taste cell. Specifically, we found that InR signaling in sated conditions dampens sweet GRN cellular responses to reduce taste sensitivity and feeding. These findings broaden our understanding of gustatory-metabolic integration and demonstrate how endocrine signaling directly alters chemosensory input to modify behavior.

## RESULTS

### Transcriptomic analysis of candidate modulatory receptors in sweet GRNs

To explore how internal metabolic state may impact peripheral taste processing, we first searched through available transcript databases. A microarray dataset from the proboscis and maxillary palps of fed and starved flies (GEO: GSE27927) revealed that *InR* and *AKHR* levels fluctuated significantly in a state-dependent manner: in flies starved one or two days, *InR* levels increased (4.06 and 5.55 fold change, respectively) and *AKHR* levels decreased (-4.27 and - 5.35 fold change, respectively)^77^. These reciprocal changes in *InR* and *AKHR* expression could potentially reflect transcriptional feedback mechanisms that help maintain homeostatic signaling during transitions in satiety state as Dilp and AKH levels rapidly fluctuate^68,75,78–80^. To search specifically for hunger/satiety hormone receptor expression in sweet GRNs, we used the single-cell transcriptomic database, Fly Cell Atlas^81^. Putative sweet GRNs were identified via expression of *Gr64f*, a sugar-specific gustatory receptor (GR) and marker of each sweet GRN on the labellum^31,46,47^. 33 total *Gr64f+* cells were isolated from a database cluster previously annotated “Gustatory receptor neuron of the labellum” (E-MTAB-10519) (Fig. 1A, Fig. S1). Within this subset, we screened for receptors with the potential to modulate sweet GRN activity.

**Figure 1:**
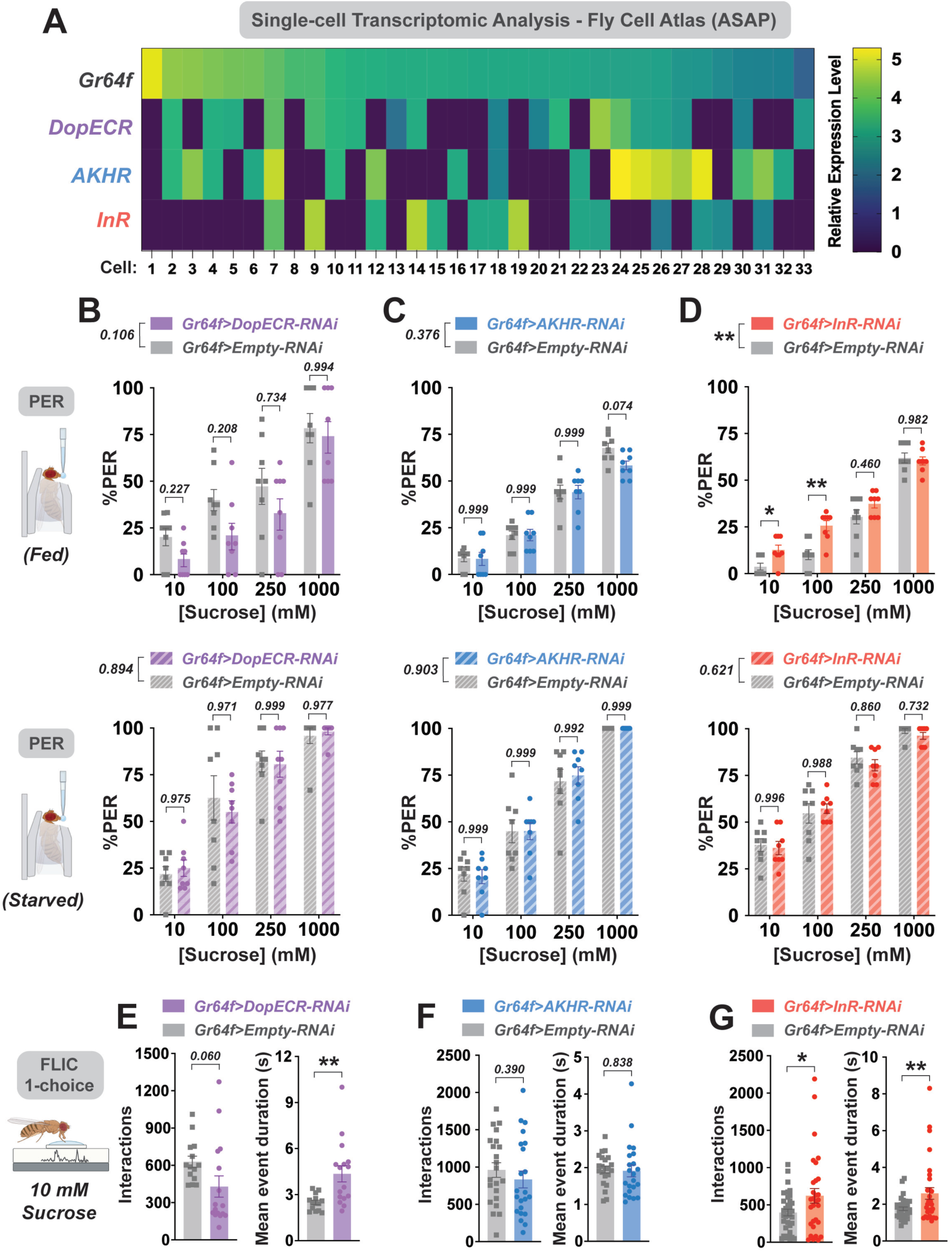
Insulin receptor knockdown in sweet GRNs affects sucrose sensitivity and feeding. (A) Expression heatmap of *Gr64f*+ sweet GRNs on the labellum and several candidate modulatory receptors via single-cell transcriptomic analysis from Fly Cell Atlas^81^. Cells ordered from highest to lowest relative *Gr64f* expression level. Warmer colors represent increased relative expression values. (B-D) Labellar PER with a sucrose concentration series (10-1000 mM) in flies expressing RNAi against *DopECR* (B), *AKHR* (C), *InR* (D), or empty RNAi controls in sweet GRNs under both fed (upper panels) and 24-hour starved (lower panels) conditions. *n* = 8 groups of 6-10 flies per genotype. All mated females. All data plotted as mean ± SEM. *p* values represent main effects of genotype by two-way ANOVA. Multiple comparisons by Šidák’s post-test at each individual concentration are depicted. *\*p < 0.05, **p < 0.01*. (E-G) 1-choice FLIC (fly liquid interaction counter) assay with 10 mM sucrose to quantify feeding interactions (left panels) and mean feeding event duration (right panels). Data from fed flies expressing RNAi against *DopECR* (E), *AKHR* (F), *InR* (G), or empty RNAi controls in sweet GRNs. *n* = 13-30 flies per genotype. All mated females. All data plotted as mean ± SEM. *\*p < 0.05, **p < 0.01* by unpaired t-test.

As a positive control, we first searched for expression of *DopECR*, a receptor previously shown to be expressed in sweet GRNs and mediate dopamine’s hunger-dependent enhancement of sweet taste sensitivity^56^. Consistent with previous results, we found *DopECR* expression in the majority of *Gr64f*+ cells (Fig. 1A, Fig. S1B). Expanding our search to metabolic receptors, *AKHR* (55%) and *InR* (42%) showed the next highest levels of expression in this subset (Fig. 1A, Fig. S1B), a finding that aligns with a previous study that used TRAP-seq of *Gr5a*+ sweet GRNs to detect a high abundance of active *InR* mRNA^82,83^. Conversely, we found no expression of two receptors previously linked to sweet GRN modulation: *CCKLR-17D3*^61^ and *NPFR*^84^ (Fig. S1A). In contrast to mammalian work demonstrating a role for leptin signaling in TRCs^85^, we did not find expression of the fly leptin receptor analog: *dome* (Fig. S1A). Despite the apparent prevalance of other neurotransmitter receptors in sweet GRNs (Fig. S1A), we decided to focus our subsequent analyses on InR and AKHR, as both endocrine receptors exhibit high *Gr64f+* co-expression and have been strongly associated with hunger/satiety regulation^67–70^. We additional analyses using other sugar-sensing GRs to demonstrate that expression of our candidate receptors is not enriched in any particular subset of sweet GRNs (Fig. S2A). Moreover, we found very low expression of *InR* and *AKHR* in *Gr66a*+ bitter GRNs^40,86^, suggesting their expression is sweet-biased and not common to all GRN populations (Fig. S2B).

### Insulin receptors expressed in sweet cells impact sucrose taste sensitivity and feeding

We were next interested in how these candidate modulatory receptors modify sweet taste sensitivity in the context of hunger or satiety. As a comparison to previous findings, we first selectively knocked-down *DopECR* via targeted RNAi only in *Gr64f*+ cells. We assessed how this manipulation altered taste sensitivity to sucrose on the labellum via the proboscis extension response (PER) assay^87^ under both fed and 24-hour starved conditions. In the sated state, *DopECR-RNAi* flies displayed lower PER rates than *Empty-RNAi* controls on average, however these differences were statistically insignificant (Fig. 1B, upper panel). Under 24-hour starved conditions, *DopECR* knockdown did not change sucrose sensitivity (Fig. 1B, lower panel), replicating previous results^56^. We then repeated this approach for both *AKHR* and *InR* knockdown. *AKHR* knockdown had no effect on sucrose sensitivity, regardless of internal state (Fig. 1C). However, *InR* knockdown resulted in a significant enhancement of sucrose sensitivity in the fed state, particularly at low concentrations (Fig. 1D). These results imply that *InR* in sweet GRNs helps to regulate sucrose sensitivity, particularly under sated conditions.

While PER assesses acute taste detection, we were also interested in how sweet GRN-specific candidate receptor knockdown would impact overall feeding behavior. To quantify nuanced feeding metrics, we utilized the Fly Liquid Interaction Counter (FLIC)^34,88^. Fed flies were individually placed into feeding chambers and given access to a 10 mM sucrose food source for 3 hours (1-choice FLIC assay). Food interactions, feeding events, and average feeding event durations were quantified for each fly. *DopECR* knockdown in sweet GRNs resulted in a trending suppression of sucrose interactions (Fig. 1E) and a strong reduction in feeding events (Fig. S1C). Interestingly, these flies also had a significant increase in the average feeding event duration (Fig. 1E), potentially suggesting the involvement of compensatory post-ingestive mechanisms that lengthen feeding bouts and maintain sufficient caloric intake during this assay. In contrast to the strong effects of *DopECR* knockdown, flies expressing *AKHR-RNAi* did not exhibt shifts in any of the tested FLIC feeding metrics (Fig. 1F, S1D). Aligning with our observed PER phenotype, *InR* knockdown in sweet GRNs also significantly elevated feeding interactions and increased average feeding event duration (Fig. 1E), but did not impact the number of feeding events (Fig. S1E). Importantly, recent work comparing FLIC and other feeding assays showed that feeding bout duration is strongly correlated with food intake^89^. Together, these findings indicate that InR’s impacts on sweet GRNs and sucrose sensitivity also meaningfully affect prolonged feeding behavior.

Since *InR* knockdown in sweet GRNs resulted in robust behavioral phenotypes compared to the other candidate receptors, we decided to focus our subsequent analysis on the impacts of InR-mediated modulation. Accordingly, we next validated *InR* expression in sweet GRNs by co-staining labellar tissue from *Gr64f>GFP* flies with a human insulin receptor (*INSR*) antibody previously shown to successfully label *InR* in larval olfactory neurons^90^. There was prominent immunoreactivity throughout the labellum, including clear overlap with *GFP*+ sweet GRN cell bodies in the sensilla and along the labial nerve (Fig. S3A). This fluoresence was absent in control tissue without the primary antibody (Fig. S3B). This finding is consistent with our analysis of the labellar transcriptome and provides protein-level confirmation for InR’s presence in sweet GRNs.

### Insulin receptor signaling in sweet cells modulates taste sensitivity and sucrose feeding in a state-dependent manner

To thoroughly investigate how InR signaling influences sweet GRN sensitivity, we expressed either a dominant-negative (*InR[DN]*) or constitutively-active (*InR[CA]*) insulin receptor transgene in *Gr64f*+ cells to induce inactive or overactive insulin signaling, respectively^68,90^. We first tested how expression of *InR[DN]* in sweet GRNs shifted sucrose sensitivity in mated female flies using sucrose PER under both fed and 24-hr starved conditions. We hypothesized that inactive InR signaling would have the strongest effects on taste sensitivity in the fed state, when circulating insulin is relatively high^91^. Indeed, sated *InR[DN]* flies had significantly elevated PER rates compared to *InR[WT]* controls, at all tested sucrose concentrations (Fig. 2A, upper panel). Under starvation conditions, PER rates between *InR[WT]* and *InR[DN]* were not statistically significant (Fig. 2A, lower panel).

**Figure 2:**
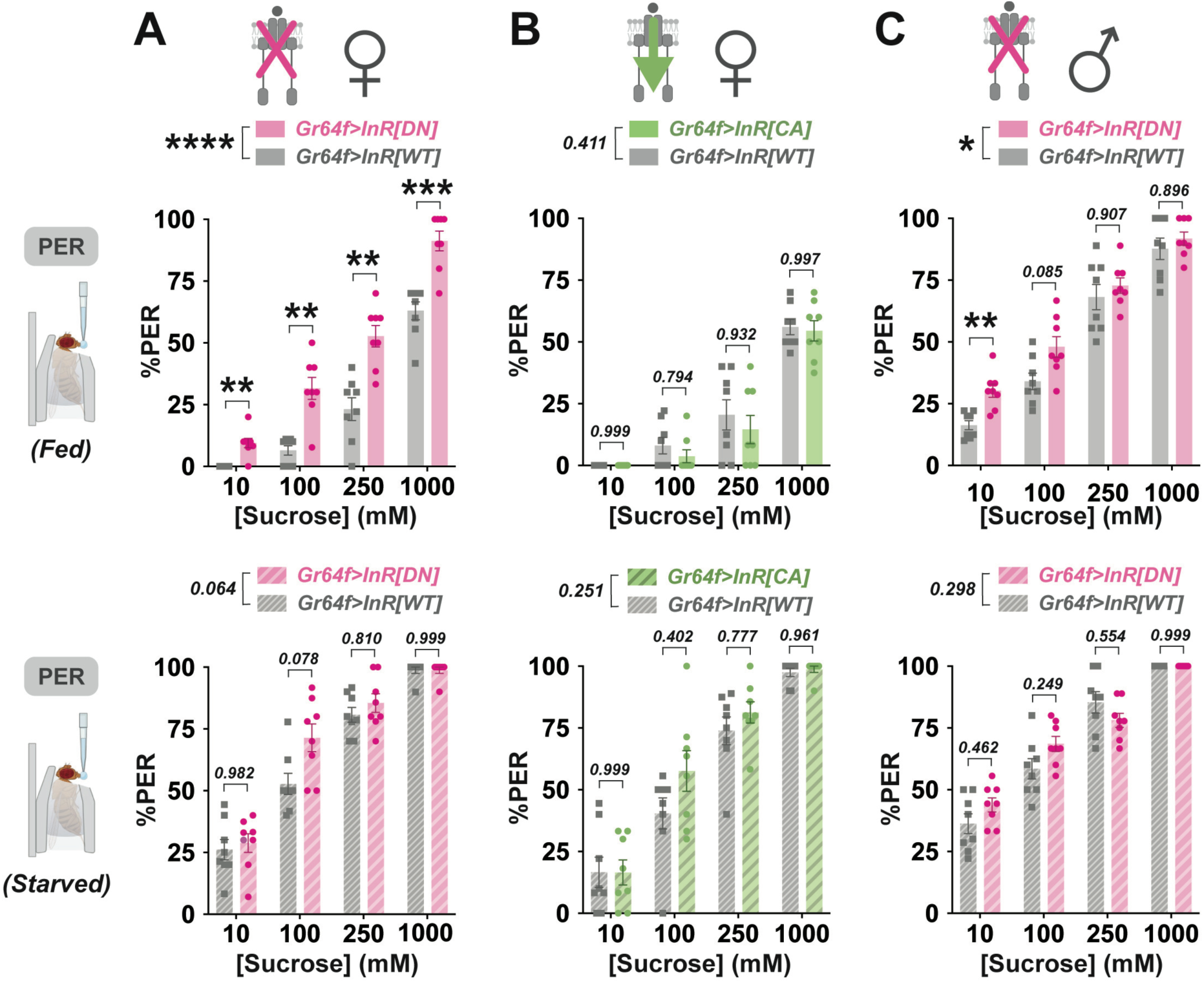
Inactive insulin receptor signaling in sweet GRNs enhances sucrose sensitivity. (A-C) Labellar PER to a sucrose concentration series (10-1000 mM) in mated female (A, B) and male (C) flies expressing dominant negative (*DN*) (A, C), constitutively active (*CA*) (B), or wildtype (*WT*) *InR* in sweet GRNs. Experiments conducted under both fed (upper panels) and 24-hour starved (lower panels) conditions. *n* = 8 groups of 9-14 flies per genotype. All data plotted as mean ± SEM. *p* values represent main genotype effects by two-way ANOVA. Multiple comparisons by Šidák’s post-test at each individual concentration are depicted. *\*\*p < 0.01, ***p < 0.001*.

Next, we predicted that overactive insulin signaling via *InR[CA]* expression would most likely reduce PER in the starved state, when Dilp signaling is relatively low^67^. However, we found no differences between *InR[CA]* flies and *InR[WT]* controls in either the fed or starved experiment (Fig. 2B). We also tested whether the behavioral impacts of overactive InR signaling would manifest when flies were moderately hungry instead of fully starved, however sucrose PER in flies food-deprived for 6 hours was also unaffected by *InR[CA]* expression (Fig. S5A). Due to well-documented sex differences in *Drosophila* insulin sensitivity and signaling^92–94^, we also modified InR signaling in male flies and assessed changes in sucrose sensitivity using the same PER paradigm. Like females, *InR[DN]* expression in male flies resulted in higher sucrose PER, especially at lower concentrations, but only in the fed state (Fig. 2C). Importantly, *Gr64f* is only expressed in adulthood^46^, thus the *Gr64f-gal4* driver should not be active throughout development to have indirect effects on adult flies. To directly determine if prolonged loss of InR signaling in adult sweet GRNs elicits gross metabolic dysregulation that could otherwise confound our behavioral phenotypes, we quantified starvation sensitivity and found that *InR[DN]* expression did not impact starvation resistance for either female or male flies (Fig. S4A). To determine if inactive InR signaling also affects sweet GRNs in the tarsi^95,96^, we first identified *Gr64f*+ sweet GRNs in the leg-specific dataset from Fly Cell Atlas and searched for *InR* co-expression. While fewer tarsal cells exhibited *Gr64f* expression, the proportion that co-express *InR* was similar to that observed in the labellum (Fig. S4B). Accordingly, we repeated our sucrose PER experiments but instead stimulated flies’ legs. Tarsal PER rates were not affected by *InR[DN]* expression in either the fed or starved state (Fig. S4C) suggesting InR’s modulatory effects are specific to labellar taste cells. Together, these results demonstrate that inactive InR signaling in labellar sweet GRNs, for both sexes, causes a fed-state specific elevation in sucrose PER, suggesting that InR typically acts to temper sweet sensitivity with satiety.

We were also interested in how InR signaling within sweet GRNs regulates overall feeding behavior and consumption. We first returned to our 1-choice FLIC assay to determine how inactive InR signaling in sweet GRNs would impact prolonged feeding behaviors. In mated females, *InR[DN]* and *InR[WT]* flies shared a similar number of feeding interactions and events (Fig. 3A, Fig. S4D), but *InR[DN]* flies exhibited longer feeding event durations at both 10 and 1000 mM sucrose (Fig. 3A). In males, *InR[DN]* flies had significantly more feeding interactions at several sucrose concentrations and showed longer feeding event durations at 10 mM (Fig. 3B) but no strong changes in the number of feeding events (Fig. S4E). Recent work comparing FLIC and other feeding assays showed that feeding bout duration is strongly correlated with food intake^89^. Aligning with our PER findings, overactive InR signaling via *InR[CA]* expression did not impact FLIC feeding metrics in flies pre-starved for 6 or 24 hours (Fig. S5B, S5C).

**Figure 3:**
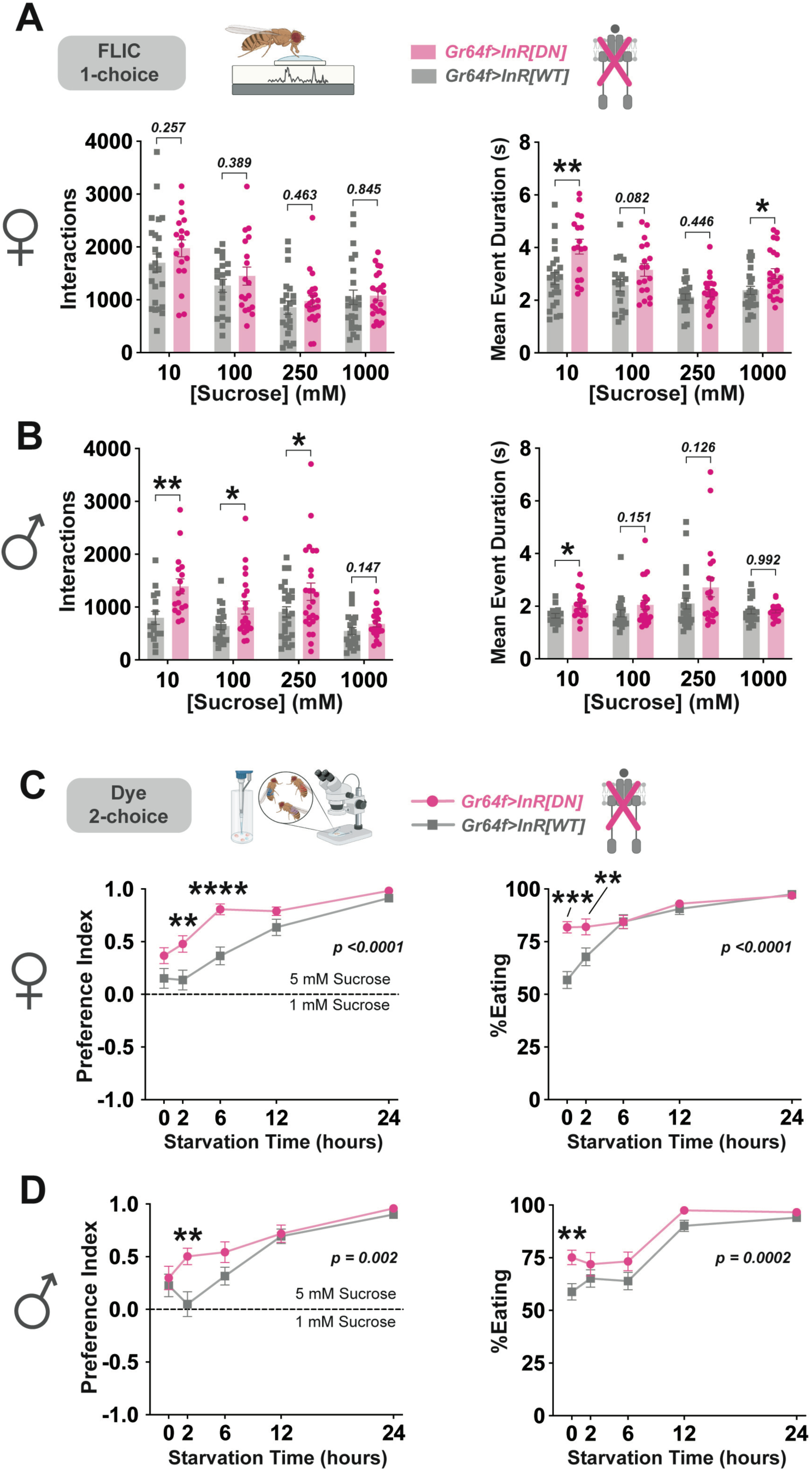
Inactive insulin signaling in sweet GRNs increases sucrose intake behaviors. (A, B) 1-choice FLIC (fly liquid interaction counter) assay at increasing sucrose concentrations (10-1000 mM) to quantify feeding interactions (left panels) and mean feeding event duration (right panels). Data from mated female (A) and male (B) flies expressing dominant negative (*DN*) or wildtype (*WT*) *InR* in sweet GRNs. *n* = 15-24 flies per condition. All data plotted as mean ± SEM. *\*p < 0.05, **p < 0.01* by unpaired t-test. (C, D) Dye-based binary feeding choice assay to measure threshold-level (1 mM vs. 5mM) sucrose preference (left panels) and consumption (right panels) at increasing starvation time points (0-24 hours). Data from mated female (C) and male (D) flies expressing dominant negative (*DN*) or wildtype (*WT*) *InR* in “sweet” GRNs. *n* = 19-21 groups of 10 flies per condition. All data plotted as mean ± SEM. *p* values represent main genotype effects by two-way ANOVA. Multiple comparisons by Šidák’s post-test at each individual starvation level are depicted. *\*\*p < 0.01, ***p < 0.001*, *\*\*\*\*p < 0.0001*.

To further characterize the behavioral effects of InR signaling, we used a dye-based binary feeding choice assay^31–33^ to measure preferential consumption of a low concentration of sucrose. For one hour, flies were allowed to freely choose between and consume two solutions: 1 mM or 5 mM sucrose, the approximate *Drosophila* threshold preference level for sucrose^97^. We conducted these experiments at five different starvation time points (0, 2, 6, 12, and 24 hours) to determine if the effect of InR would change with increasing hunger. Both female and male *InR[DN]* flies had elevated preference and consumption of this low concentration of sucrose compared to *InR[WT]* controls, particularly at low/moderate hunger levels (Fig. 3C-D). Consistent with the rest of our behavioral results, preferential consumption was not impacted by *InR[CA]* expression, regardless of starvation level (Fig. S5D). Collectively, these findings suggest that InR signaling modulates sweet GRNs to reduce sweet detection thresholds and sucrose consumption.

### Insulin signaling modifies sweet taste cell calcium responses

Previous investigation of sweet GRN regulation revealed that hunger enhances cellular sucrose sensitivity via increased calcium influx^56,57^. To determine if InR signaling regulates cellular sucrose responses in a similar manner, we performed *in vivo* calcium imaging of sweet GRNs during labellar sucrose stimulation^38^ while recording changes in GCaMP6f fluorescence^98^. Specifically, we imaged sweet GRN axon terminals in the sub-esophageal zone of the brain, a region where GRNs synapse with second-order taste neurons^45^. This technique has been used to describe nuances about *Drosophila* gustatory processing in the context of internal state modulation^2,31,56,57^. We quantified calcium responses from the SEZ under both fed and 24-hour starved conditions, recording from flies expressing wildtype, inactive, or overactive InR. Between *InR[DN]* and *InR[WT]* flies, there was no overall effect of genotype for either satiety state (Fig. 4A, S6B). However, under fed conditions, *InR[DN]* flies exhibited a significant increase in peak calcium responses to 10 mM sucrose (Fig. 4A, S6A). Conversely, constitutively active insulin signaling during starvation significantly suppressed calcium responses to sucrose. *InR[CA]* flies showed a significant decrease in peak fluorescence compared to *InR[WT]* controls, most prominently to 10 mM and 1000 mM sucrose (Fig. 4B). Expression of *InR[CA]* did not alter any cellular calcium responses in the fed state (Fig. S6C). These results demonstrate that inactive and overactive InR lead to reciprocal and state-dependent effects on sweet GRN calcium responses, further confirming the suppression of neuronal taste responses by InR, at least at lower concentrations.

**Figure 4:**
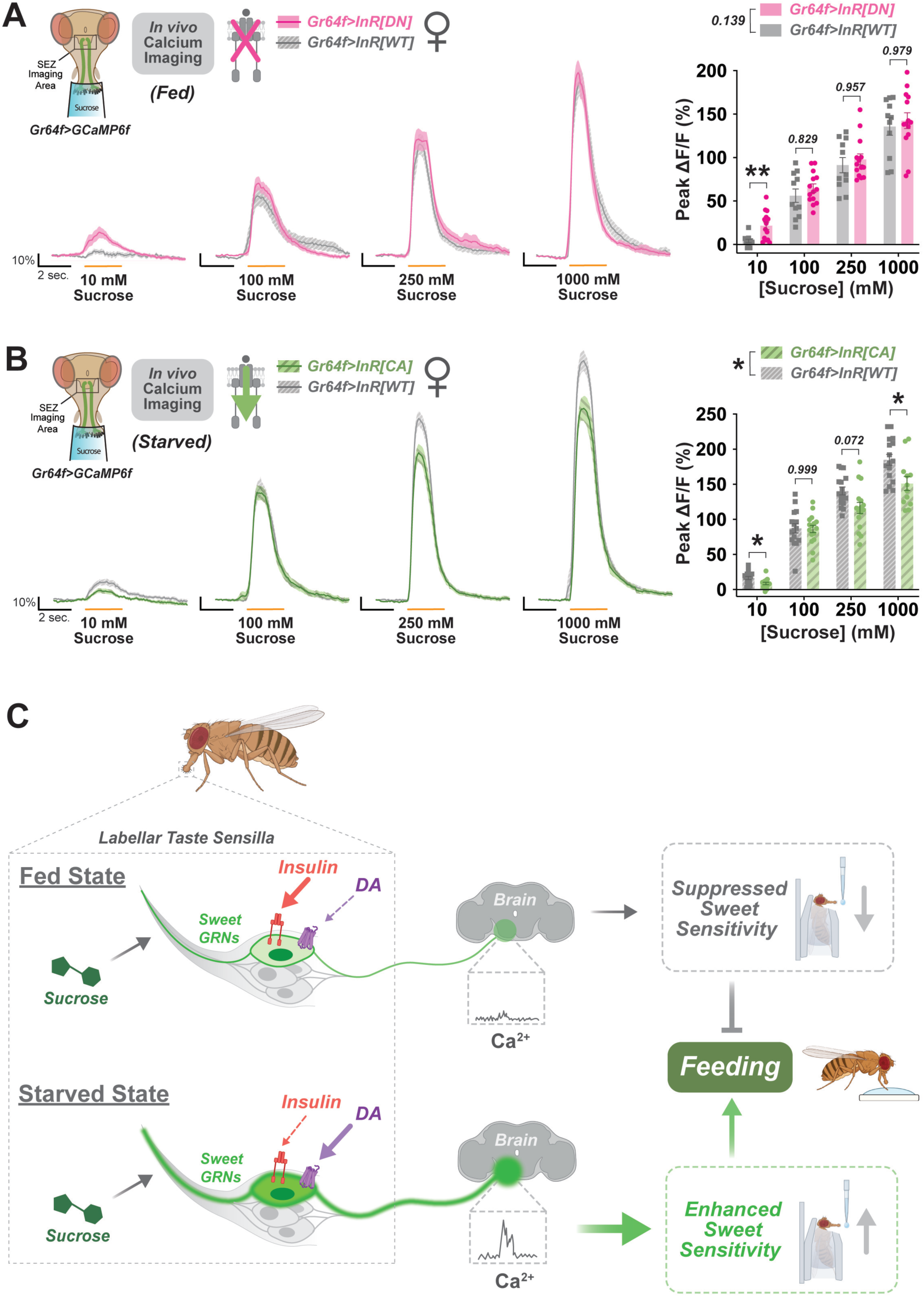
Insulin receptor signaling in sweet GRNs influences cellular responses to sucrose. (A, B) *In vivo* calcium imaging of GCaMP6f in sweet GRNs during labellar sucrose stimulation. Data from flies expressing dominant negative (*DN*) (A), constitutively active (*CA*) (B), or wildtype (*WT*) *InR* in sweet GRNs under fed (A) or 24-hour starved (B) conditions. Calcium responses measured as ΔF/F over time at each concentration (left) and peak ΔF/F (right). Yellow lines under each curve indicate when the stimulus was on the labellum. *n* = 11-16 flies per genotype. All mated females. All data plotted as mean ± SEM. *p* values represent main genotype effects by two-way ANOVA. Multiple comparisons by Šidák’s post-test at each concentration are depicted. *\*p < 0.05*. (C) Model for satiety state-dependent modulation of sweet GRNs by InR. In the fed state, InR signaling (red) in sweet GRNs dampens cellular sensitivity to sucrose and reduces feeding. In the starved state, hunger signals like dopamine (DA) via DopECR (purple) plus downstream circuit modulation can impact sucrose sensitivity and promote feeding.

## DISCUSSION

Describing the molecular and cellular basis of metabolic-gustatory interactions is essential for understanding the pathological consequences of their dysregulation. For example, sweet taste sensitivity in healthy individuals fluctuates throughout the day^99^, but this variation is absent in those with insulin resistance^100^. In addition, chemosensory disruption is a well-documented symptom of various metabolic disorders^101–103^, but we are only beginning to elucidate the impact of sensory nutrition^104^ on overall health. From our basic investigation in *Drosophila*, we propose that high circulating insulin/Dilps act on InRs expressed in sweet GRNs to suppress cellular responses to sucrose, reduce sweet sensitivity, and deter feeding in a fed state, but in a starved state, any signal from low circulating insulin/Dilps is potentially overridden by other hunger signals that enhance sweet sensitivity (Fig. 4C.) These hunger signals may include dopamine released onto sweet GRNs via DopECR^56,57^ or additional downstream modulatory mechanisms in the brain^37,105^ that increase sweet salience and encourage feeding (Fig. 4C). These results offer insight into potentially conserved mechanisms of sensory modulation by metabolic signals.

### Negative regulation of sweet taste sensitivity by insulin receptors

Mammalian investigations of gustatory regulation at the level of taste bud have implicated several neuropeptide and endocrine signals, including: neuropeptide Y^106^, cholecystokinin^107^, leptin^85^, and glucagon^65^. Insulin receptor expression has also been observed in type II TRCs^66^, but their role in taste sensitivity was not evaluated. Aligning with our single-cell transcriptomic analysis, previous studies implementing TRAP-seq of *Gr5a*+ sweet GRNs detected a high abundance of active *InR* mRNA^82,83^ and recent work using the same Fly Cell Atlas database also reported *InR* expression in primary GRNs^108^. Importantly, the internal conditions of the flies used for Fly Cell Atlas could impact expression levels, since at least *InR* and *AKHR* fluctuate with metabolic status^77^.

Previously, one study found no changes in sucrose PER when *Dilp3*-expressing cells were acutely silenced^58^, implying that insulin signaling would have limited influence on sweet taste responsiveness. However, this study used a global silencing approach and performed PER under 24-hour starved conditions. Our InR manipulation was specific to sweet GRNs, and the resulting phenotype only emerged in the fed state, when circulating Dilp levels are relatively high^91^. Furthermore, cholecystokinin-like DSK acts as a prominent satiety signal in the fly brain and is co-released by insulin-producing cells (IPCs) to reduce food intake^60^. Another study provided evidence that DSK suppresses sweet taste within GRNs following refeeding, but targeted knockdown of *CCKLR-17D3* implied that DSK receptor signaling actually had the opposite effect - enhanced sweet sensitivity^61^. In addition to the lack of expression in our Fly Cell Atlas search (Fig. S1A), this discrepancy complicates the current interpretation of DSK and CCKLR-17D3 involvement in sweet GRNs.

Prior investigation found that starvation-dependent sucrose sensitivity required DopECR in sweet GRNs, but their knockdown of *DopECR* only reduced sucrose PER following 6 hours of starvation and, consistent with our results, this effect was not seen at 24 hours^56^. Other previous studies have identified mechanisms of sweet GRN regulation by internal state in response to dietary manipulation. A high sugar diet has been shown to dampen sweet GRN sensitivity through the activation of the intracellular sugar sensor: O-linked N-Acetylglucosamine transferase (OGT)^34^ and through a gut-taste axis mediated by Hedgehog signaling^109^. Importantly, our tarsal PER results indicate that InR does not significantly modulate sweet GRNs in the legs (Fig. S4C). Previous work using tarsal PER found that prolonged exposure to dietary sucralose enhances sweet GRN sensitivity via NPF, a *Drosophila* neuropeptide Y analog, and its receptor, NPFR^84^. Thus, it is possible that modulation occurs through distinct pathways on different sensory organs. Other studies have instead proposed that NPFR more generally contributes to hunger-enhanced sweet sensitivity via upstream excitation of dopaminergic interneurons in the brain^57,58^, and we were unable to find *NPFR* expression in labellar *Gr64f+* cells (Fig. S1A). Consistent with previous studies, we found strong and specific expression of *AKHR* in sweet GRNs (Fig. 1A)^8,110^. However, we did not find any impact of this receptor on sweet sensitivity or feeding behavior (Fig. 1C, S1D). While the exact function of this glucagon-like metabolic receptor within sweet taste cells remains unclear, past work found that AKHR expressed in the brain mediates starvation-induced hyperactivity^8^. This result could suggest that AKHR signaling in sweet GRNs could contribute to other hunger-dependent behaviors.

### Modification of feeding behavior from insulin signaling in taste cells

In addition to its metabolic functions, insulin helps control energy homeostasis by signaling satiety in the brain^111^. In mammals, this regulation is characterized by anorexigenic circuitry in the hypothalamus^111^, but insulin can also interact with dopaminergic reward pathways to affect sweet valence^112^. In *Drosophila*, global *InR* inactivation in the brain leads to increased preference for caloric sugar, an effect that mimics a food-deprivation phenotype and further supports the anorexigenic nature of insulin signaling in the fly nervous system^113^. In the current study, our dye-based binary feeding choice assays showed that InR signaling in sweet cells diminishes sucrose detection thresholds and consumption. Additional experiments using FLIC revealed that these same signals shorten the duration of sucrose feeding bouts, a metric that is representative of food intake^89^. Importantly, inactive InR signaling in sweet GRNs did not impact starvation survival (Fig. S4A), suggesting these changes in feeding output were likely not driven by metabolic dysfunction. The phenotypes we observed in our dye-based assays were most prominent at low/moderate starvation levels and the FLIC results were in fed flies, further confirming the state-dependent nature of insulin signaling in sweet GRNs.

While both feeding assays have been used to uncover connections between taste processing and behavior^31–34^, these experiments measure sucrose intake over multiple hours. This raises the possibility that our feeding phenotypes were impacted by post-ingestive mechanisms within the brain^114^ or gut^115^ that regulate food intake over longer periods of time. Despite this concern, we saw significant enhancement in feeding behavior when only primary GRNs were manipulated. A potential explanation is that increased consumption caused by the loss of satiety signaling in sweet cells increases secretion of Dilps which are still unable to modulate taste responses, leading to even more consumption. Moreover, sweet GRNs in the pharynx express *Gr64f* and are known to regulate prolonged sugar feeding, suggesting these feeding phenotypes could also reflect modulation of pharyngeal sweet GRNs. Additionally, studies in humans^116^, rodent models^117^, and *Drosophila*^92–94^ highlight significant sex differences in insulin signaling and sensitivity. Despite these important differences, our data indicate that InR similarly modulates sweet GRNs in both sexes. InR inactivation significantly enhanced sucrose sensitivity (Fig. 2A, 2C), feeding behavior (Fig. 3A-B) , or preferential consumption (Fig. 3C-D) in both male and female flies. The one subtle dimorphism we observed was *InR[DN]* expression having a stronger enhancing effect on sucrose interactions in male flies compared to females of the same genotype (Fig. 3B). Along with previous evidence showing that insulin pathway activity is elevated in males on a low-sugar diet^93^ our finding could suggest that the male gustatory system is more sensitive to InR modulation under certain conditions. However, future work is necessary to fully characterize how these potential sex differences manifest at a behavioral level.

### Direct modulation of cellular taste responses by insulin receptor signaling

Since our InR manipulation impacted sucrose PER and feeding, we anticipated that cellular changes in primary taste neurons may drive this behavioral shift. Measuring changes in neuronal activity with *in vivo* calcium imaging is strongly indicative of action potential firing patterns^118^ and this approach has been used to study complex aspects of gustatory processing^31–38^. For example, previous calcium imaging analysis showed that during starvation, sucrose elicits stronger responses in sweet GRNs^56,57^. In our control flies, we replicated this starvation effect and found that food-deprivation alone enhances calcium responses in sweet GRN axon terminals (Fig. 4A). Similar approaches have also revealed that sweet GRN activity can be affected by high dietary sugar^34,35^ or parallel bitter circuitry via sweet cell-specific GABA_B_R signaling^59^. Here, we propose that InR in sweet GRNs modulates neuronal activity in a state-dependent manner, adding another mechanism by which primary taste cell responses can be dynamically regulated.

Expression of *InR[CA]* in sweet GRNs significantly suppressed cellular responses to sucrose during starvation (Fig. 4B), a finding that is consistent with our initial prediction that the effects of overactive InR signaling would most likely manifest under low Dilp conditions. However, this same manipulation did not elicit behavioral phenotypes. This discrepancy could reflect redundancy in hunger signals acting downstream to preserve feeding drive even when sweet input is diminished. For example, modulatory DopECR signaling in sweet GRNs does not fully explain hunger-induced enhancement of sweet taste sensitivity^56,57^, and higher-order gustatory neurons within the PER or related circuits are also directly modulated by starvation^37,105^.

In contrast, expression of *InR[DN]* in sweet GRNs strongly enhanced sucrose PER and feeding in the fed state, but these behavioral phenotypes were associated with an increase in neuronal calcium responses only at 10 mM sucrose (Fig. 4A). This concentration likely lies near the sensory threshold for sweet GRN activation, particularly under fed conditions, and subtle elevations in calcium influx caused by inactive InR signaling may lift suppression just enough to generate a detectable increase in fluorescence. At higher sucrose concentrations, when *InR[WT]* sweet GRNs are already strongly activated, our assay may lack the sensitivity required to detect further modulatory increases in calcium signal, masking potential physiologically relevant changes in sweet GRN activity that may contribute to behavioral modification. Alternatively, mild differences in calcium responses could also indicate the prevalence of other mechanisms underlying InR-mediated sweet GRN suppression that would not be associated with changes in calcium transients, including potential rapid structural changes. In *Drosophila*, acute presynaptic homeostatic potentiation (PHP) in phasic motor neurons can occur within just 10 minutes and is not associated with changes in calcium influx, but has been shown to promote homeostatic signaling by rapidly adjusting presynaptic active zone nanostructure^119,120^. In mice, leptin induces presynaptic structural changes within hypothalamic feeding circuits after only 6 hours^121^. Moreover, evidence from other invertebrate model systems has revealed that insulin and other neuromodulators can drive presynaptic remodeling in sensory neurons after 12-24 hours^122,123^. Future investigation is required to explore the precise intracellular or circuit-level mechanisms in which InR signaling regulates sweet GRN output and the resulting behavioral consequences.

In conclusion, we describe a mechanism of direct communication between endocrine signaling and taste processing that regulates sweet sensitivity at a primary cellular and behavioral level. Future work can explore the intracellular or circuit-level mechanisms by which insulin elicits taste suppression and how higher-order taste neurons may also integrate internal state signaling to further control feeding.

### Limitations of this Study

While quantifying *InR* expression or signaling via qPCR or Western Blot would help confirm the effects of InR knockdown or inactivation, detecting these minor changes would be extremely technically challenging as we are altering InR in only one cell type across a complex sensory organ (labellum). Despite the effects elicited by our InR manipulations being consistent, we acknowledge that the magnitude of these changes were minor, especially at the cellular level, but discussion above regarding potential explanations for the discrepenacies between our behavioral and cellular findings).

## ACKNOWEDGEMENTS

We thank the Bloomington Stock Center (BDSC) for fly stocks. *UAS-InR[WT], UAS-InR[DN], and UAS-InR[CA]* were provided by Exelxis, Inc. to BDSC. We thank Dr. Dennis Matthew at the University of Nevada, Reno for the *INSR* antibody recommendation and evidence for its successful cross-reactivity in *Drosophila*. We also thank Michael Gordon and Viktoriya Li in the Gordon Lab at the University of British Columbia where pilot studies were conducted. Graphics were generated with BioRender.com and we thank Kayla Audette for contributing to the graphics. This work was funded by new lab startup funds from the University of Vermont.

## AUTHOR CONTRIBUTIONS

Conceptualization, M.S.; methodology, C.A. and M.S.; investigation, C.A., L.E., M.I, and M.S.; writing - original draft, C.A. and M.S.; writing - review & editing, C.A., L.E., M.I., and M.S.; visualization, C.A. and M.S.; resources, M.S.; supervision, M.S.; funding acquisition, M.S.

## DECLARATION OF INTERESTS

The authors declare no competing interests.

## DATA AVAILABILITY

A Mendeley dataset containing the original data generated in this study is available, DOI: 10.17632/b3y8dtwcz3.1.

## METHODS

### Flies

All flies were kept on standard cornmeal food at 25°C and 60% humidity before testing. For all experiments, flies were 2-10 days old. Mated females and males were used where indicated. Information for each genotype can be found in the key resources table. All fly stocks are available from the BDSC. Stocks were not back-crossed, rather all comparisons were made with genetic background-matched controls. Two separate *UAS-Empty-RNAi* lines were used to generate the control genotypes used in Figure 1. To match their respective experimental alleles, BDSC #36303 (chromosome 3) was used for the *DopECR* knockdown experiment and BDSC #36304 (chromosome 2) was used for the *AKHR* and *InR* knockdown experiments.

### Chemicals

A full list of chemicals with source information can be found in the key resources table. Sucrose was made up in water at 1 M and diluted further in water to the indicated concentrations.

### Fly Cell Atlas analysis

Using ASAP (https://asap.epfl.ch/fca), we analyzed the database’s 10x proboscis and maxillary palps (stringent) or 10x leg (stringent) datasets (E-MTAB-10519). We first identified the previously annotated cell clusters “gustatory receptor neuron of the labellum” from the proboscis and “gustatory receptor neuron” from the leg dataset via HVG-UMAP visualization^81^. A subset of cells in each cluster that exhibited expression of *Gr64f* were identified as sweet GRNs. In each putative sweet GRN, relative expression values for *Gr64f* and each candidate receptor were recorded from ASAP. Raw UMI reads for each cell were normalized (10,000 counts/cell) and log-transformed (log(x+1)) to generate relative expression levels for the genes of interest^81^.

### PER

Proboscis extension response (PER)^87^ was used to assess appetitive feeding responses and taste sensitivity by quantifying the proportion of proboscis extensions in response to test stimulations. Flies were collected into vials containing standard cornmeal food (fed experiments) or into food deprivation vials containing 1% agar (starved experiments) 24 hours prior to testing and kept at 25°C. For labellar PER, flies were individually mounted under a dissection scope using a mouth pipette to transfer each fly into a 200 uL pipette tip that was cut so only the head and labellum were exposed. For tarsal PER, flies were mounted on slides using quick-dry nail polish to immobilize the flies while exposing their tarsi. For all PER experiments, flies recovered in a humidity chamber for ∼1 hour after mounting and then manually water satiated. Water also served as the first stimulus to ensure subsequent PER responses were not driven by thirst or dehydration. The test solutions consisted of increasing sucrose concentrations, including 10 mM, 100 mM, 250 mM, and 1000 mM sucrose. Labellar sensilla or tarsi were manually stimulated three times for each concentration. A positive response was recorded if the fly extended to at least one stimulation. Water stimulations in between each sucrose solution were used to verify continued water satiety.

### Immunohistochemistry

Immunofluorescence on labella was conducted as previously described^31,33^. Labella from female, *Gr64f>CsChrimson.mVenus* flies (shorthand: *Gr64f>GFP)* were dissected, split, and fixed in 4% paraformaldehyde in PBS +0.2% Triton (PBST) for 25 minutes before washing.

Tissues were then blocked in 5% normal goat serum (NGS) for 1 hour before adding primary antibodies (chick anti-GFP at 1:1000, rabbit anti-InR at 1:40) overnight. A human polyclonal insulin receptor (INSR) antibody (PAA895Hu02) was chosen due to its immunogen aligning closely with the conserved region of *Drosophila InR* and previous evidence showing reactivity to InR protein in larval olfactory neurons^90^. The anti-InR primary antibody was excluded for negative control tissue, but the rest of the protocol remained the same. After washing with 0.2% PBST, secondary antibodies (goat anti-chicken Alexa 488, goat anti-rabbit Alexa 647, both at 1:200) were incubated for 3 hours. After washing in PBST, samples were placed on slides in SlowFade gold with #1 coverslips as spacers. Images were acquired with a 3i Spinning disc Confocal station (Zeiss upright microscope, 2 K × 2 K 40 fps sCMOS camera, CSU-W1 T1 50 μm spinning disc) with a 40x water immersion objective. A z-stack with 1 micron steps were captured with the following settings: laser power 40, 488 channel 20 ms exposure, 647 channel 100 ms exposure, brightfield 100 ms exposure. FIJI (ImageJ) was used to z-project 20 microns of labellar tissue (maximum projection for fluorescent channels, minimum projection for brightfield), and images were exported separately and merged with the 647 INSR blue channel pseudocolored to red for clearer visualization of overlap. All scale bars= 50 microns.

### Starvation Survival Assay

For starvation survival experiments, groups of 10 mated female or male flies were transferred into food deprivation vials (1% agar) at 2 days old and kept at 25°C for the duration of the experiment. Every 12 hours, the number of flies that were alive in each vial was recorded until all the flies had died. Female and male vials were run in parallel.

### FLIC

FLIC experiments were performed as previously described^33,88^. For each experiment, flies were taken directly off standard cornmeal food and individually transferred into behavioral feeding chambers via mouth pipette. Capacitance sensors in each chamber measured the number of interactions between a fly and a single sucrose food source (1-choice assay). Flies from each genotype were loaded into chambers on opposite sides of a *Drosophila* Feeding Monitor (DFM) and interactions with the sucrose solution were recorded over 3 hours. This raw output was analyzed via a custom R script modified from the Pletcher Lab (full code on GitHub, see Key Resources Table). Measurements from the loading process were excluded to avoid potential artifacts caused by adding flies into the chambers. Data were analyzed with R code like previous publications to quantify feeding interactions, feeding events, and feeding event durations. Minimum signal threshold was set to 10, feeding threshold was set to 20, and measurements were taken every 200ms. A feeding interaction was defined as any individual reading above the feeding threshold. A feeding event was defined as 10 consecutive interactions above the minimum threshold, as long as one of these readings reached the feeding threshold. Sequential periods of continuous interactions were combined into one single event as long as the gaps of inactivity between them lasted less than 1 second. Feeding event duration was measured in seconds. Data from chambers that exhibited disrupted signal detection were removed, including flies with 0 recorded interactions or raw values that were exceedingly high (> 1000 discrepancy between raw interaction count and R analysis output). Flies that failed to interact with the food source (<15 interactions) or exhibited extremely high mean event duration measurements (> 12 seconds) were also excluded.

### Dye-based Feeding Assays

Binary choice experiments were conducted as previously described^31–33^. Vials containing 10 flies were flipped into 1% agar food-deprivation vials for the indicated number of hours before test. Flies assessed at 0-hour starvation were flipped into vials containing standard cornmeal food. At test, flies were transferred into vials that contained six dots of alternating red (0.5 mg/mL Amaranth, FD and C Red#2) and blue (0.125 mg/mL Erioglaucine, FD and C Blue#1) colored solutions. Each 10uL dot contained dye, 2% agar, and either 1 mM sucrose or 5 mM sucrose. The color associated with each sucrose concentration was counterbalanced so that half of the replicates had red 5 mM sucrose and half had blue 5 mM sucrose. Flies were allowed to freely choose between either food option for 1 hour in the dark in an incubator with ∼50% humidity at 29°C. After being frozen at -20°C, a dissection microscope was used to score each fly’s post-mortem abdomen color as red, blue, purple, or no color. Preference index for each vial was calculated as: ((# of flies labeled with X color)-(# of flies labeled with Y color))/(total # of flies with color). The number of flies in each vial that consumed either food choice (%Eating) was calculated as: ((# of flies labeled blue, red, or purple/total # flies in vial) *100).

### *In vivo* Calcium Imaging

GCaMP6f fluorescence of Gr64f GRN terminals was performed as previously described^31,32,38^. Flies were briefly anesthetized with CO_2_ and mounted in a custom imaging chamber with the proboscis extended and held in place with a dental waxer. Mounted flies recovered for one hour in a humidity chamber, then a small area of cuticle was removed and the esophagus cut to expose the SEZ. Adult hemolymph-like (AHL) solution (108 mM NaCl, 5 mM KCl, 4 mM NaHCO_3_, 1 mM NaH2PO_4_, 5 mM HEPES, 15 mM ribose, 2 mM Ca^2+^, 8.2 mM Mg^2+^, pH 7.5) was continuously applied to the area during the dissection and the same solution was used for immersion during imaging. A Leica SP5 II Confocal microscope with a 25x immersion objective captured the GCaMP6f signal over time. The SEZ was imaged at 4-6x zoom with an 8000Hz line speed, line accumulation of 2, and a resolution of 512 x 512 pixels with the pinhole opened to 2.86 AU. A micromanipulator with a glass capillary delivered the solutions (water or sucrose) over the labellum while the GCaMP6f signal was recorded. Each stimulation included 5 seconds of baseline, 2 seconds with the stimulus over the labellum, and 7 seconds while the signal returned to baseline after stimulus removal. Each fly received water first, followed by increasing concentrations of sucrose. Fed flies were kept on food while starved flies were kept on 1% agar for 24 hours prior to mounting. Experiments comparing fed and starved flies of the indicated genotypes were run in parallel on the same day. FIJI (ImageJ) was used to draw a tight ROI around sweet GRN axonal projections in the SEZ via baseline GCaMP6f signal^38^. Flourescence values from the ROI was then extracted over time. For each GCaMP6f recording, ten timepoints in a row were used as a baseline and the signal converted to ι1F/F (%). The peak ι1F/F was calculated as the maximum change in fluorescence using an average of three timepoints around the highest value.

### Statistical Analysis

All statistical analyses were conducted using GraphPad Prism 10 software. Sample sizes were chosen based off estimated effect sizes and power analyses from previous experiments using similar assays. Genotype controls were run in parallel for each experiment. Specific statistical tests are included in the figure legends. FLIC experiments were conducted separately at each sucrose concentration and genotypes were compared using unpaired t-tests. For all other experiments, sucrose concentrations or starvation time points were run in parallel and genotype differences were quantified by two-way ANOVA (genotype main-effect) with Šidák’s post-test. All data plotted as mean ± SEM. Asterisks indicate *\*p < 0.05, **p < 0.01, ***p. < 0.001, ****p < 0.0001*.

## Key Resources Table

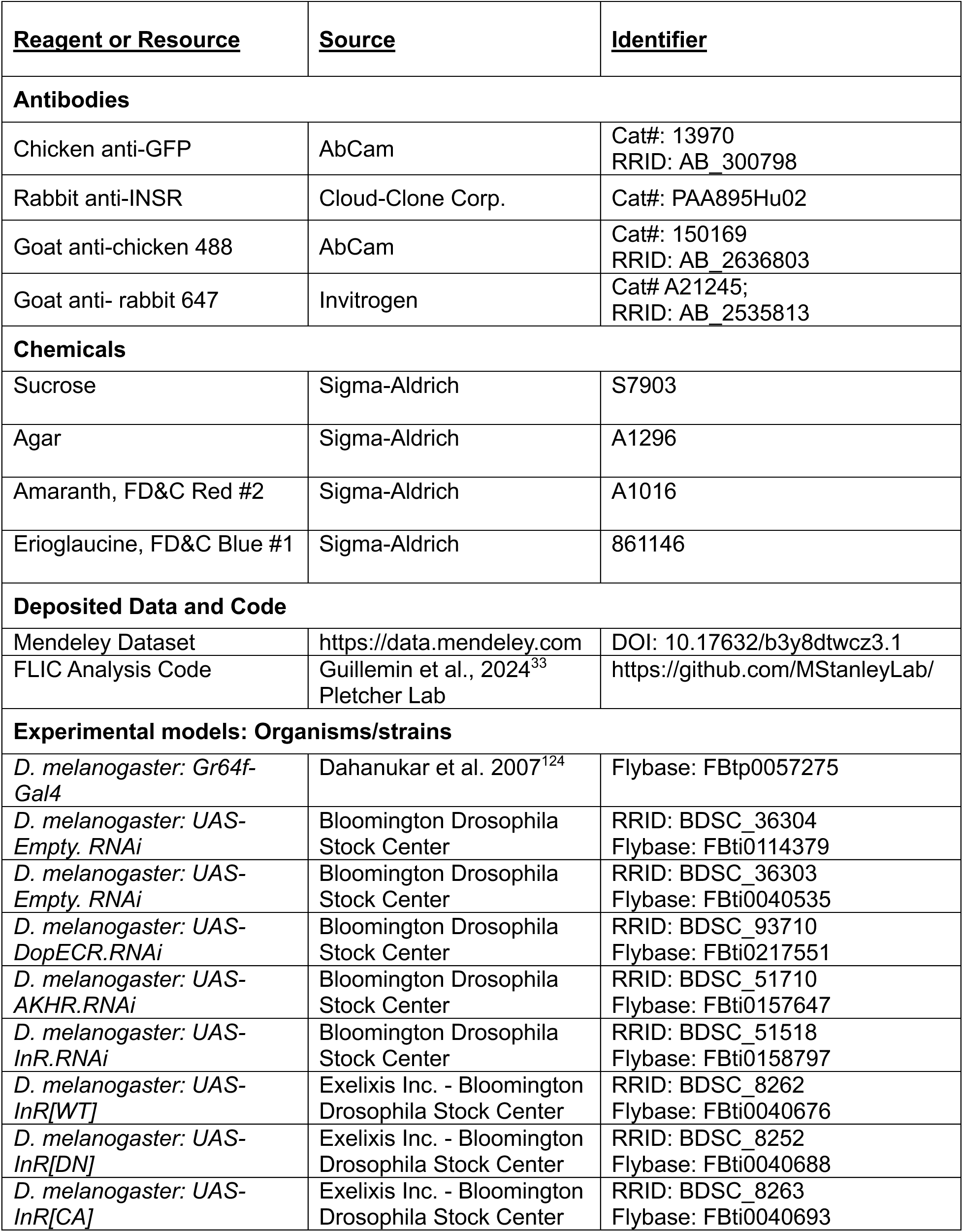
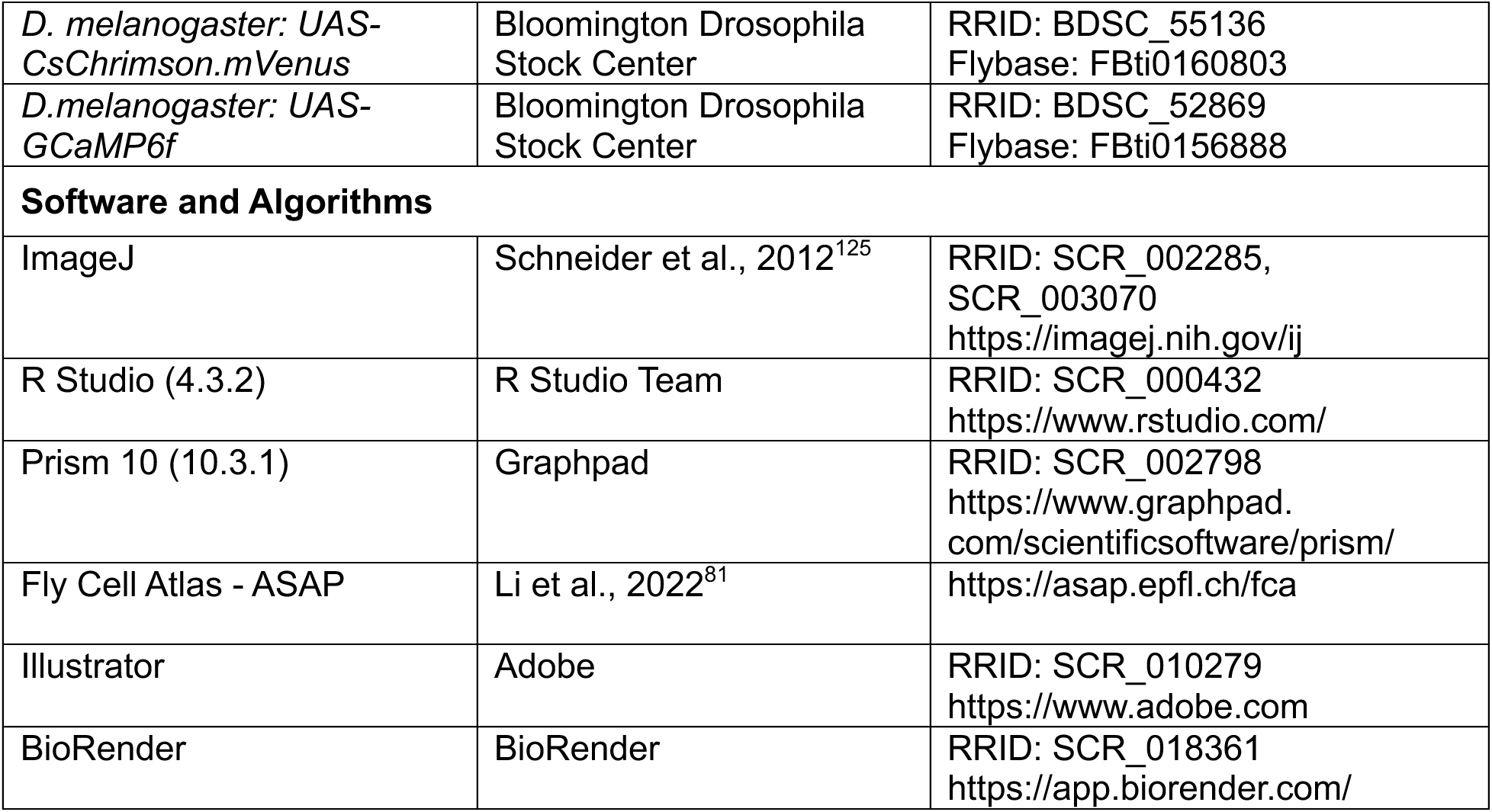

## SUPPLEMENTAL

**Figure S1:**
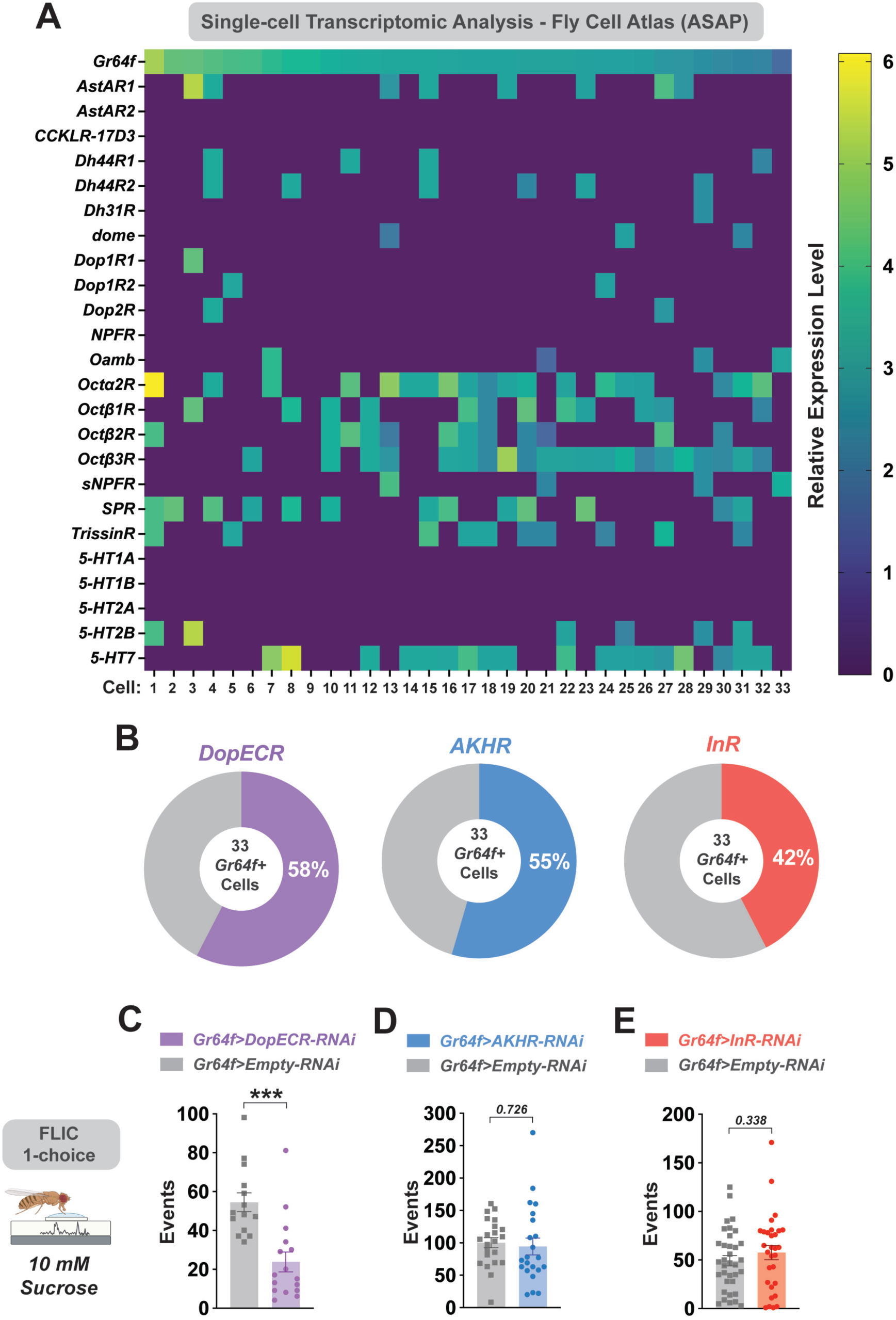
Sweet GRNs co-express combinations of candidate modulatory receptors and only DopECR knockdown impacts feeding events. (A) Expression heatmap of all *Gr64f+* sweet GRNs showing a panel of other candidate modulatory receptors. (B) Breakdown of all *Gr64f*+ sweet GRNs by expression of three candidate receptors from Figure 1A: *DopECR* (purple), *AKHR* (blue), and *InR* (red). All data from single-cell transcriptomic analysis with Fly Cell Atlas – ASAP. (C-E) 1-choice FLIC (fly liquid interaction counter) assay with 10 mM sucrose to quantify feeding events, accompanying data from Figure 1E-G. Data from fed flies expressing RNAi against *DopECR* (C), *AKHR* (D), *InR* (E), or empty RNAi controls in sweet GRNs. *n* = 13-30 flies per genotype. All mated females. All data plotted as mean ± SEM. *\*\*\*p < 0.001* by unpaired t-test.

**Figure S2:**
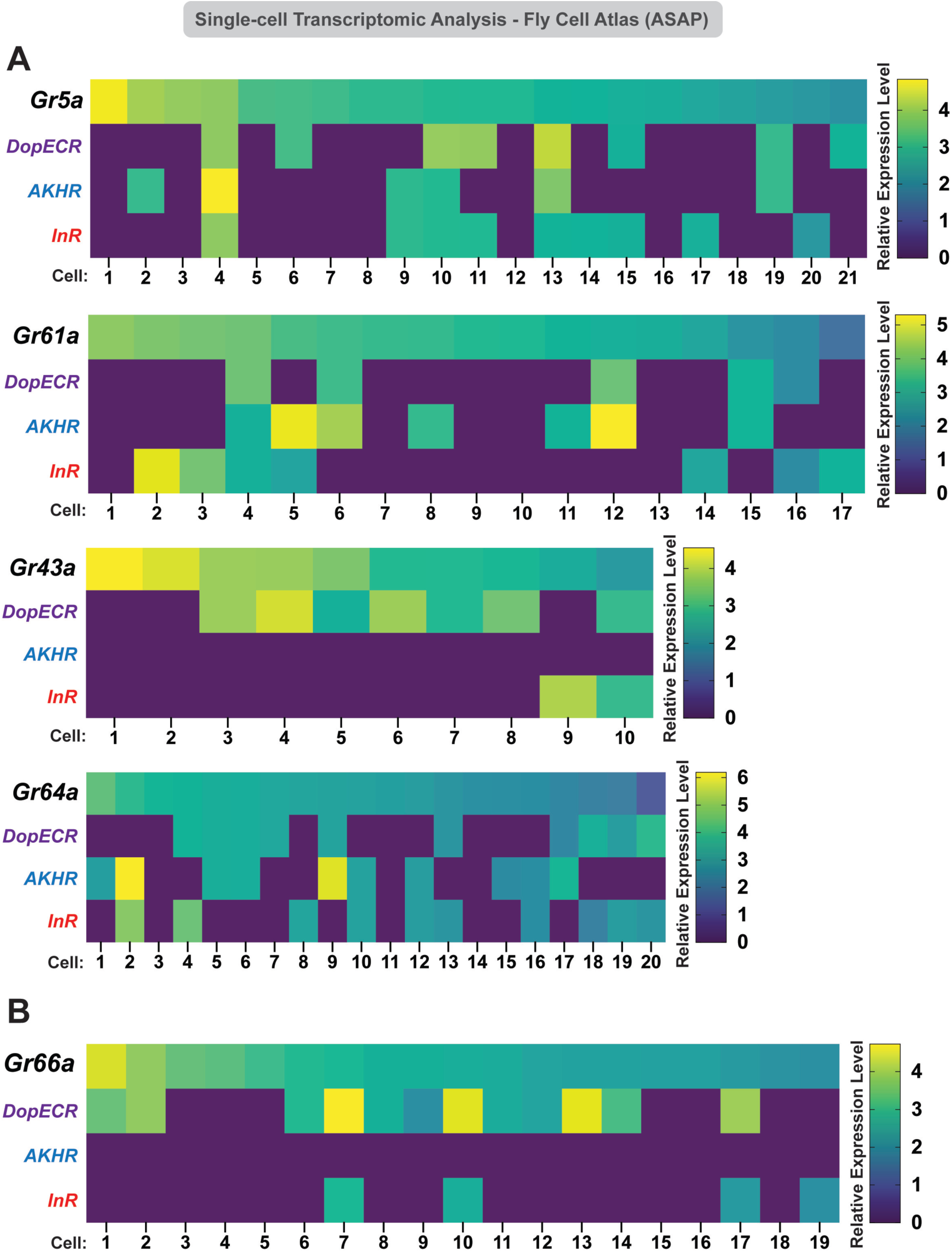
Modulatory receptor expression is similar in sweet GRN subpopulations and low in bitter GRNs. (A,B) Expression heatmap of sweet GRNs labeled by alternative sugar GRs (A) and bitter GRNs labeled by *Gr66a* (B) showing primary candidate modulatory receptors. All data from single-cell transcriptomic analysis with Fly Cell Atlas – ASAP.

**Figure S3:**
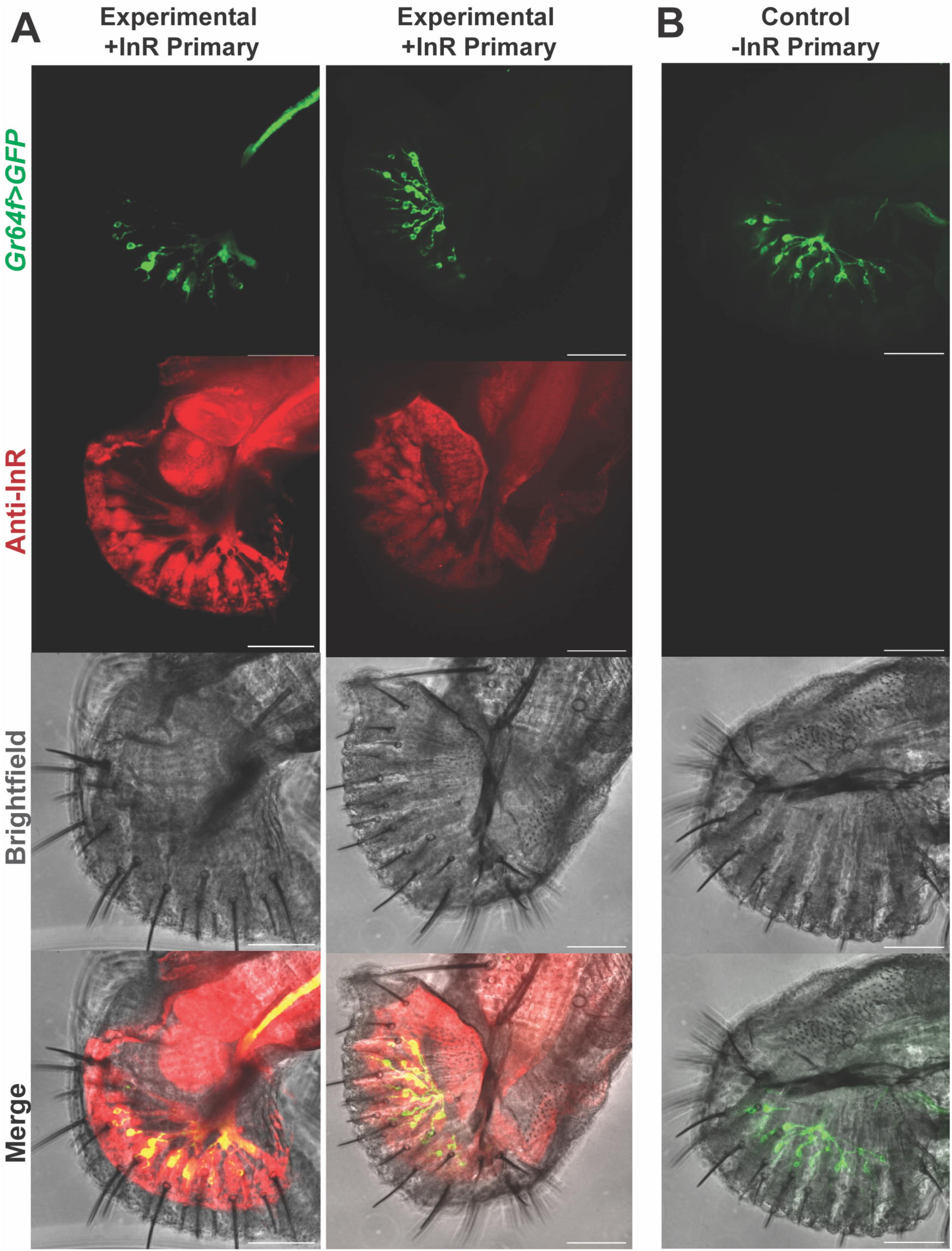
Immunohistochemical validation for InR expression in sweet GRNs. (A,B) Split labellar palps immunolabelled for *Gr64f-gal4* driving *UAS-CsChrimson.mVenus* (shorthand: *Gr64f>GFP*) with InR co-staining using a human *INSR* antibody. Representative images from two experimental labella that were exposed to the primary INSR antibody (A) and from a negative control without the primary antibody (B) are shown. *Gr64f>GFP* (green), *anti-InR* (red), and brightfield channels are depicted separately (top 3 rows) and merged together (bottom row). Yellow color in GRN cell bodies and along labial nerve represent overlapping *Gr64f>GFP* and InR signal. Tissue obtained from mated females. Scale bars= 50 µm.

**Figure S4:**
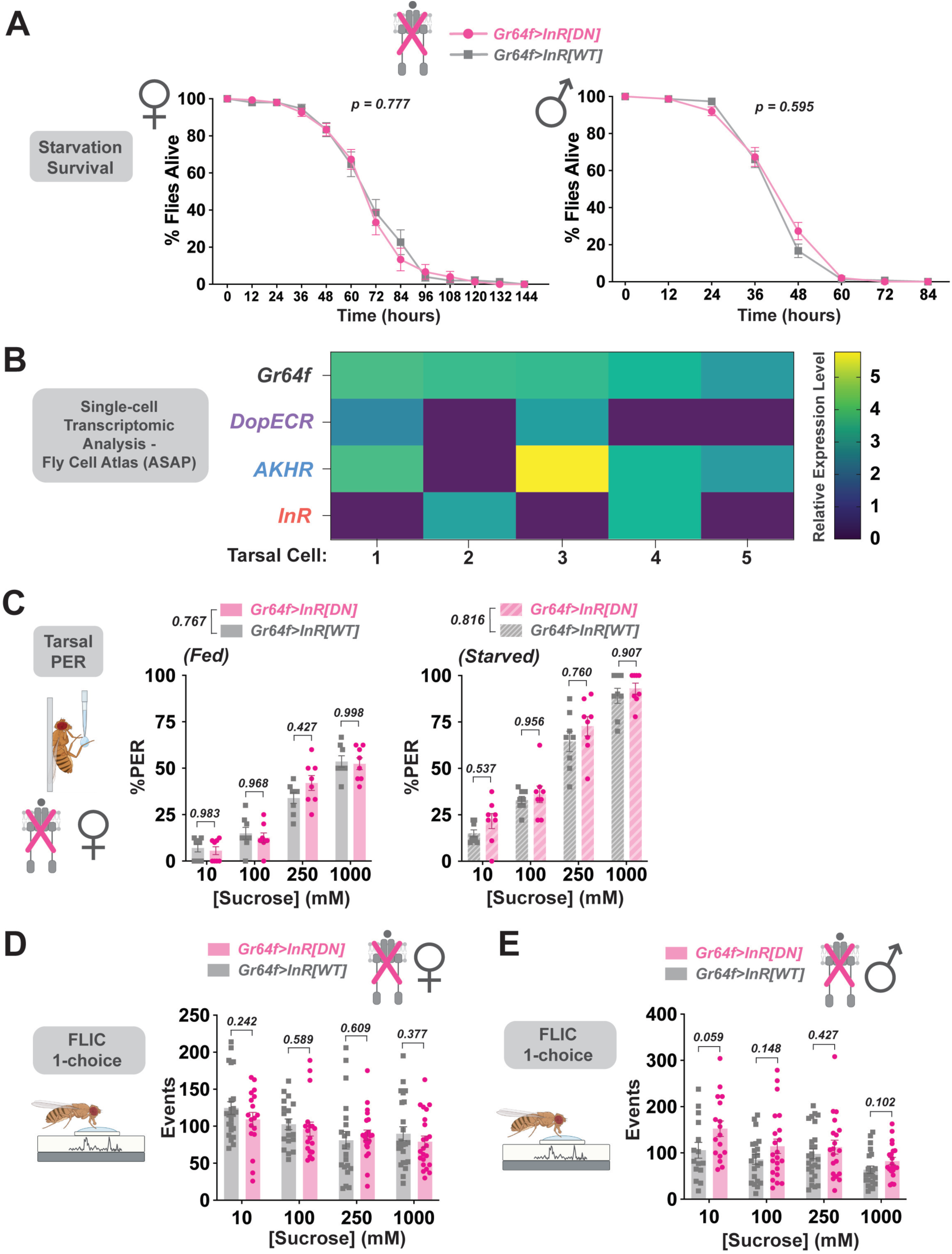
InR inactivation does not impact starvation sensitivity, tarsal PER, or feeding events. (A) Starvation survival assay to assess gross metabolic dysfunction in both mated females (left panel) and males (right panel). *n* = 15 vials of 10 flies per genotype. Data plotted as mean ± SEM %flies alive after each 12-hour timepoint. Main genotype effects by two-way ANOVA with Šidák’s post-test are depicted. (B) Expression heatmap of *Gr64f*+ sweet GRNs in the legs and several candidate modulatory receptors via single-cell transcriptomic analysis from Fly Cell Atlas. Cells ordered from highest to lowest relative *Gr64f* expression level. Warmer colors represent increased relative expression values. (C) Tarsal PER to a sucrose concentration series (10-1000 mM) in mated female flies expressing wildtype (WT) or dominant negative (*DN*) *InR* in sweet GRNs. Experiments conducted under both fed (left panel) and 24-hour starved (right panel) conditions. *n* = 8 groups of 7-10 flies per genotype. Data plotted as mean ± SEM. *p* values represent main genotype effects by two-way ANOVA with Šidák’s post-test. (D,E) 1-choice FLIC (fly liquid interaction counter) assay at increasing sucrose concentrations (10-1000 mM) to quantify feeding events in mated female (D) and male (E) flies expressing dominant negative (*DN*) or wildtype (*WT*) *InR* in sweet GRNs. *n* = 15-24 flies per condition. All data plotted as mean ± SEM. *p* values via unpaired t-test are shown.

**Figure S5:**
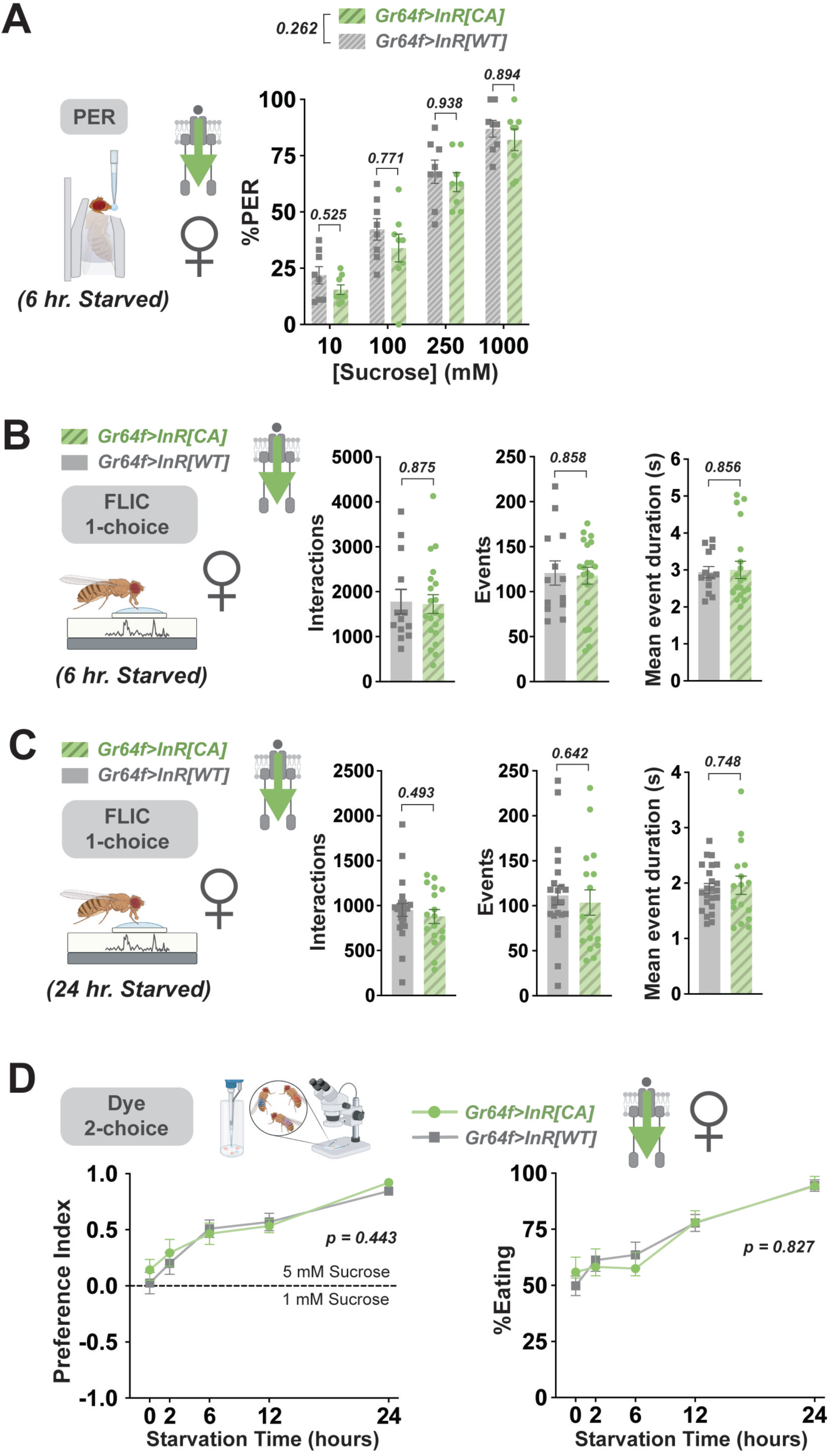
Overactive InR signaling has no impact on sucrose PER or feeding. (A) Labellar PER with a sucrose concentration series (10-1000 mM) in flies expressing constitutively active (CA) or wildtype (WT) InR in sweet GRNs. Experiments conducted under 6 hr.-starved conditions. n = 8 groups of 7-10 female flies per genotype. All data plotted as mean ± SEM. *p* values represent main genotype effects by two-way ANOVA and comparisions at each individual concentration via Šidák’s post-test. (B, C) 1-choice FLIC (fly liquid interaction counter) assay at with 10 mM sucrose to quantify feeding metrics in flies expressing constitutively active (*CA*) or wildtype (*WT*) *InR* in sweet GRNs. Both genotypes pre-starved for 6 (B) or 24 (C) hours. *n* = 13-23 flies per genotype. All data plotted as mean ± SEM. *p* values via unpaired t-test are shown. (D) Dye-based binary feeding choice assay to measure threshold-level (1 mM vs. 5 mM) sucrose preference (left panel) and consumption (right panel) at increasing starvation time points (0-24 hours). Female flies expressing constitutively active (*CA*) or wildtype (*WT*) *InR* in sweet GRNs. *n* = 20-28 groups of 10 flies per condition. All data plotted as mean ± SEM. *p* values represent main genotype effects by two-way ANOVA.

**Figure S6:**
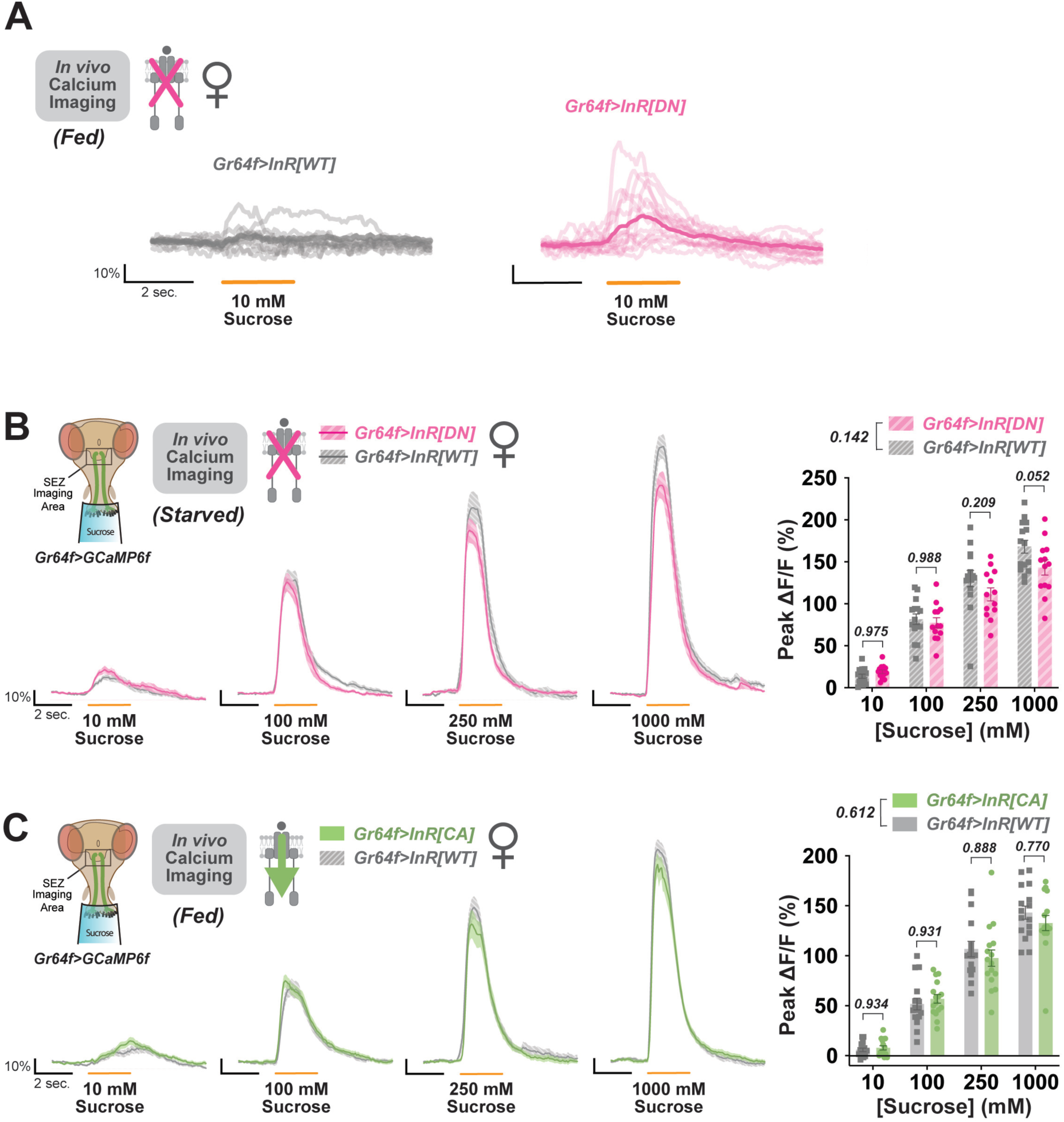
Minimal cellular effects of inactive insulin signaling in the starved condition or of overactive signaling in the fed condition. (A - C) *In vivo* calcium imaging of GCamp6f in sweet GRNs during labellar sucrose stimulation. Data from flies expressing dominant negative (*DN*) (A, B), constitutively active (*CA*) (B), or wildtype (*WT*) *InR* in sweet GRNs under fed (A, C) or 24-hour starved (B) conditions. Calcium responses measured as ΔF/F over time at each concentration for all individual replicates (A) or as ΔF/F over time at each concentration (right) and peak ΔF/F (left) as mean ± SEM (B, C). All mated females. Main genotype effects by two-way ANOVA and Šidák’s post-tests are depicted.

## Notes

### Competing Interest Statement

The authors have declared no competing interest.

